# Weberian interval discrimination and temporal pattern generalization in *Drosophila*

**DOI:** 10.64898/2026.02.04.703894

**Authors:** Tsz-Nga Lo, Chen-Ying Huang, Suewei Lin

## Abstract

Recognizing temporally structured signals, from human speech and music to the songs and calls of other animals, relies on comparing the durations of their elements. In vertebrates, interval discrimination often follows Weber’s law: sensitivity depends on the ratio between intervals rather than their absolute difference. Such proportional scaling may enable efficient representation of a wide range of durations and support recognition of temporal patterns despite changes in their speed. Whether insect brains use the same computational principle, and which neural circuits support flexible temporal learning, remain unclear. Here, we developed an auditory learning paradigm in which fruit flies, *Drosophila melanogaster*, learn to associate temporal patterns of sound with a sugar reward. Flies discriminate tone durations ranging from subsecond to a few seconds and use this information to guide reward-seeking behavior. Their discrimination depends on the ratio between durations rather than their absolute difference, consistent with Weber’s law. Flies also generalize learned temporal relationships to proportionally compressed durations and transfer them from auditory to visual stimuli, indicating a flexible representation of temporal structure. Finally, blocking mushroom body output impairs temporal pattern learning but leaves auditory pitch learning intact, identifying the mushroom body as a critical neural circuit for this form of learning. Together, these findings demonstrate that ratio-based and generalizable temporal computations can be implemented in a compact, genetically tractable brain, providing insight into the evolutionary origins of temporal cognition and establishing *Drosophila* as a powerful system for investigating its neural basis.

## Introduction

Temporal cognition—the ability to measure, compare and generalize temporal relationships—is a fundamental computation of nervous systems. Many natural signals carry meaning not only through their physical features, but through the order, duration, and spacing of their elements. Spoken words, musical rhythms, birdsong, and insect courtship displays all depend on such temporal structure. To interpret these signals, animals must extract temporal relationships from continuous sensory experience and use them to guide behavior. Because temporally structured signals are widespread across the animal kingdom, studying how brains of different sizes, architectures, and ecological demands process temporal information offers a window into whether nervous systems share conserved computational principles for timing. Such comparisons can also shed light on the neural mechanisms and evolutionary origins of temporal cognition.

At the core of temporal cognition is interval comparison. Each element of a temporal pattern contributes not only through its absolute duration, but also through its duration relative to the other elements in the sequence. In humans and many other vertebrates, interval discrimination is often governed not by the absolute difference between two durations, but by their ratio^1–9^. This proportional scaling is consistent with Weber’s law, a fundamental principle of perception in which the discrimination threshold increases with stimulus magnitude^10,11^. From an information-theoretic perspective, proportional scaling is important in its own right because it provides an efficient coding strategy: by representing larger intervals with progressively lower resolution than shorter ones, a nervous system with finite coding capacity can represent a substantially wider range of durations while preserving behaviorally relevant relative differences^12,13^.

Furthermore, Weberian interval discrimination suggests that temporal patterns may be represented in terms of their relative, rather than absolute, durations. Such a ratio-based representation could support two forms of temporal generalization. First, it could allow animals to recognize the same temporal pattern from the relative temporal relationships when it unfolds faster or slower, and the absolute durations no longer match those previously experienced. For example, human speech accelerates with emotion, a predator’s footsteps speed up when it runs, and an insect’s courtship song changes with temperature^14^. Recognizing these signals despite such natural variation would provide a clear adaptive advantage. Second, a ratio-based representation also suggests a transformation from raw sensory-specific temporal inputs into a more abstract internal representation of time. Such an abstract representation could allow animals to recognize the same temporal relationships regardless of whether they are conveyed by sound, vision, or another sensory modality. Despite the fundamental importance of Weberian timing as a principle of interval comparison, it remains unclear whether such relative and abstract temporal representations extend beyond vertebrates as general features of temporal cognition across diverse animal brains.

Studies in invertebrates suggest that compact nervous systems can also process temporal information to guide behavior across diverse ecological and experimental contexts^15–21^. However, most of these behaviors reflect timed responses that are tuned to species-specific temporal patterns or expressed at specific intervals on which the animals were trained. These stimulus-bound behaviors do not reveal whether these animals can compare time intervals or use temporal information in a flexible manner. Recently, independent of our work, two findings suggest the possibility of comparison-based temporal cognition in bumblebees. Bumblebees can discriminate the durations of light stimuli^22^ and generalize learned rhythms across tempi and transfer them across sensory modalities^23^. Together, these findings demonstrate that insect brains can support flexible temporal cognition. However, they also raise a more fundamental question: what computational principle governs interval comparison? In vertebrates, interval discrimination follows Weber’s law. Yet whether insect interval discrimination obeys the same principle remains unknown. Resolving this question would establish the computational principle of interval comparison and provide a computational framework for investigating its neural implementation, which remains difficult to dissect in bees because of the lack of genetic tools. Equally important is the question of how broadly this capacity is shared. Because bumblebees are eusocial insects with rich social and foraging ecologies, it remains unclear whether flexible temporal cognition is a specialized adaptation to these demands or a more fundamental property shared by even non-eusocial nervous systems.

The fruit fly *Drosophila melanogaster* provides an ideal system for addressing these conceptual questions. As a non-eusocial insect with a compact brain of approximately 140,000 neurons^24^, *Drosophila* offers a stringent test of whether flexible temporal cognition depends on large or socially complex nervous systems. At the same time, its extensive genetic toolkit enables circuit-level dissection of the neural mechanisms underlying temporal computation. Here we developed a classical conditioning paradigm in which flies learn to use temporally patterned stimuli to guide reward-seeking behavior. Using this approach, we asked what principle governs interval discrimination and, in particular, whether it is consistent with Weber’s law. We then asked how flexibly temporal relationships are represented across changes in duration and sensory modality, and whether the mushroom body (MB), a major associative learning center in the fly brain, contributes to temporal pattern learning.

## Results

### Flies can discriminate different stimulus intervals

To begin investigating temporal cognition in *Drosophila*, we first asked whether flies can discriminate stimulus intervals and use this information to guide behavior. We developed an associative learning paradigm inspired by classical olfactory conditioning using a T-maze^25^.

In traditional reward-based olfactory learning assays, flies learn to associate a sugar reward with one of two training odors and will subsequently approach the rewarded odor when given a choice. A prerequisite for these assays is that flies must be able to distinguish between these two odors. Here, we adapted this framework by replacing odor cues with auditory stimuli that differed in their temporal structure.

The training protocol consisted of two distinct phases (Steps 1–4 in Fig. 1a). In the first phase (Steps 1 and 2), flies were exposed to two tone patterns and learned to associate one of them (CS+) with a sugar reward. The second phase (Steps 3 and 4) then reinforced this initial association by presenting the two tone patterns in alternation from opposite ends of the T-maze. This alternating-cue training established a directional relationship between the tone patterns and reward, such that approaching the CS+ tone source led to sugar. It also familiarized the flies with the stimulus configuration used in the subsequent test phase (Step 5 in Fig. 1a).

**Figure 1:**
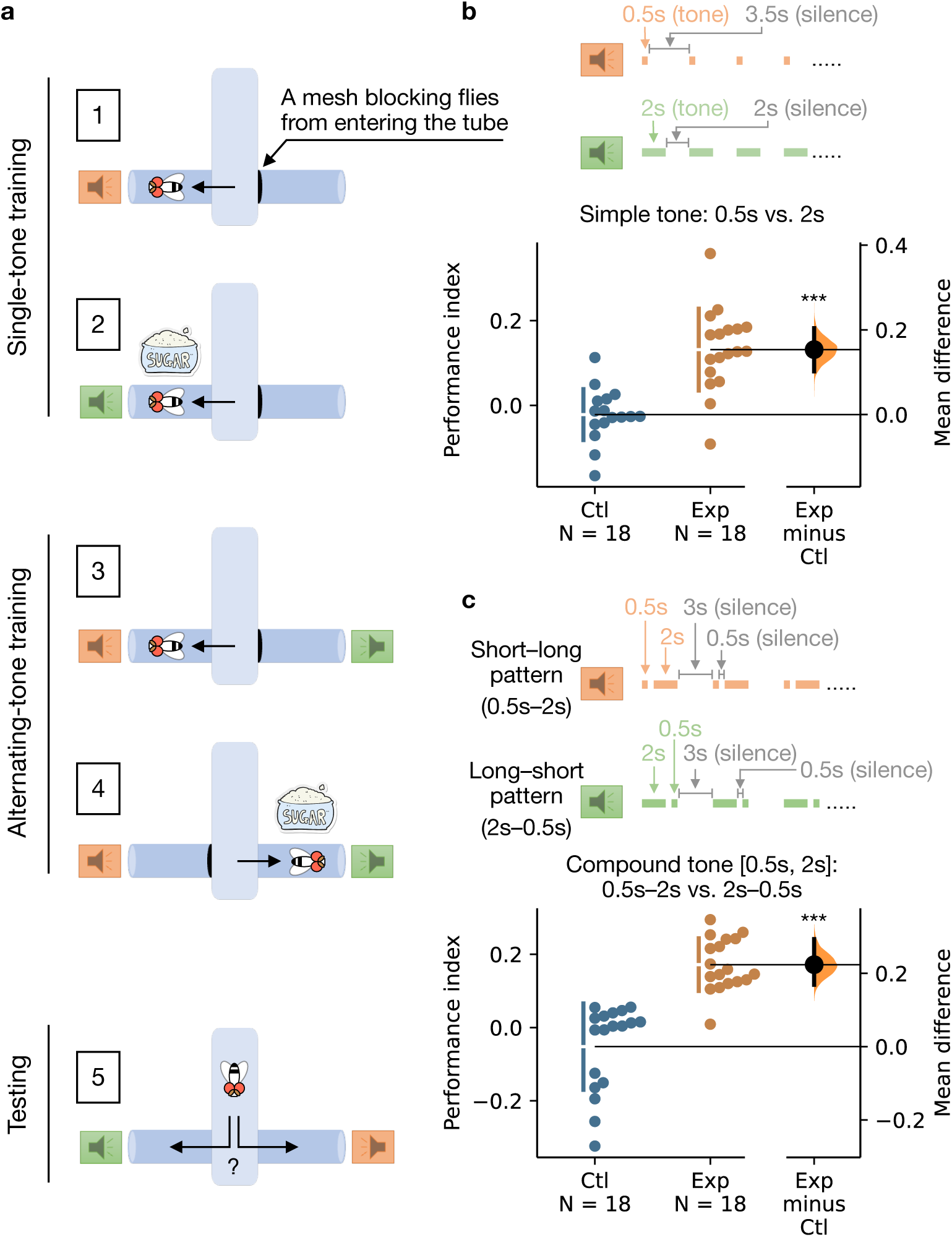
Flies learn to use tone-duration patterns to guide reward-seeking behavior. (a) Schematic of the training paradigm for associating a sugar reward with tone patterns of different durations. Flies starved for 24 h were exposed to two tone patterns composed of different tone durations. One pattern (CS–) was presented without sugar (Step 1), and the other (CS+) was paired with sugar (Step 2), each for 2 min. The flies were then presented with both tone patterns in alternation: first near the CS– pattern (no sugar) for 2 min (Step 3), then near the CS+ pattern (with sugar) for 2 min (Step 4). This alternating-cue training reinforced the CS+ association and established a directional relationship (i.e., approaching the CS+ tone source led to reward). The sides where the tone patterns were played were then swapped, and the flies were immediately tested for their preference between CS+ and CS– while both tone patterns were played in alternation (Step 5). To control for innate preferences, the tone identities were counterbalanced as CS+, and performance scores were calculated as the average of two reciprocal training groups. (b) Flies preferred the tone pattern previously associated with sugar. Simple tone patterns consisted of either repeated 0.5-s tones with 3.5-s silences or 2-s tones with 2-s silences. In the experimental group (Exp), sugar was paired with CS+ tones; in the control group (Ctl), flies underwent the same procedure without sugar. This definition of Exp and Ctl groups was used in all subsequent figures. (c) Same as (b), but with compound tone patterns composed of short and long tones. The short–long sequence included a 0.5-s tone, 0.5-s silence, 2-s tone, and 3-s silence; the long–short pattern used the same structure but with the tone order reversed. The square bracket notation (e.g., [0.5 s, 2 s] in this figure) denotes the pair of tone durations forming each compound pattern, with both temporal orders (short–long and long–short) presented during training and testing. The estimation plots in (b) and (c) show mean differences between Exp and Ctl groups, with bootstrap 95% confidence intervals shown on the right. *P*-values were calculated using Monte Carlo permutation test: ^∗∗∗^*P* < 0.001.

We first designed two simple tone patterns as the auditory cues, composed of repeated tones of either 0.5-second (s) or 2-s duration, both played at 150 Hz and 90 dB (Fig. 1b). As in olfactory learning, successful task performance required flies to discriminate between these two tone patterns. Flies were first presented with one tone pattern (CS–) for 2 minutes (min) without sugar (Step 1), followed by exposure to the other tone pattern (CS+) paired with sugar, also for 2 min (Step 2). Next, the two tone patterns were presented in alternation, with one cycle of CS– followed by one cycle of CS+, repeated continuously. The CS– and CS+ patterns were delivered from opposite ends of the T-maze. Flies were placed first in the arm near the CS– source without sugar for 2 min (Step 3), and then in the arm near the CS+ source with sugar for another 2 min (Step 4).

After training, flies were given a choice between the CS+ and CS– tone patterns for 2 min. To ensure that their decisions were based on the tone patterns rather than the spatial location of the arms, we swapped the positions of the tone patterns during testing. To control for any innate preference for one tone pattern over the other, we calculated a performance index by averaging the results from two groups of flies: one trained with the short-duration (0.5 s) tone pattern as CS+, and the other trained with the long-duration (2 s) tone pattern as CS+.

Trained flies exhibited a stronger preference for the CS+ tone pattern compared to control flies, which underwent the same training procedure but without the sugar reward (Fig. 1b). Since the main difference between the CS+ and CS– patterns was the duration of the tone, this result suggests that flies can distinguish between 0.5-s and 2-s tone durations and use this temporal information to guide foraging decisions. However, a potential concern with this interpretation is that the long-tone pattern contains more total sound than the short-tone pattern. Thus, flies may have distinguished the patterns based on total sound amount rather than on temporal structure.

To address this concern, we designed two new compound tone patterns, each consisting of a repeated cycle of short- and long-duration tones (Fig. 1c). In the short–long pattern, each cycle included a 0.5-s tone followed by a 2-s tone; in the long–short pattern, the order was reversed. These two patterns had the same total tone durations but still tests whether flies can distinguish between tone durations of 0.5 s and 2 s. If flies could not distinguish these two durations, they would perceive both patterns as identical and thus wouldn’t be able to make a correct choice during testing. Flies trained with these patterns exhibited a stronger preference for the CS+ pattern during testing (Fig. 1c), confirming their ability to distinguish between 0.5-s and 2-s tone durations. Based on these results and to eliminate differences in total sound amount, we used compound tone patterns in all subsequent experiments.

### Temporal discrimination in flies follows Weber’s law

Interval discrimination in humans and many other vertebrates has been shown to follow Weber’s law, which states that the smallest detectable change in a stimulus is proportional to the magnitude of the original stimulus—meaning that temporal perception is sensitive to ratios rather than absolute differences^3–9^. If interval discrimination in flies also follows Weber’s law, it would suggest that ratio-based temporal processing can be implemented in compact invertebrate brains and may reflect a conserved or convergently evolved computational strategy for time perception.

To test this possibility, we trained flies using compound tone patterns composed of different short- and long-duration tone pairs. As stated earlier, flies must perceive the duration difference between the short and long tones to be successfully trained. We first kept the difference between the short and long durations (ΔTime) constant, and varied the ratio between them by increasing or decreasing both durations (Fig. 2a).

**Figure 2:**
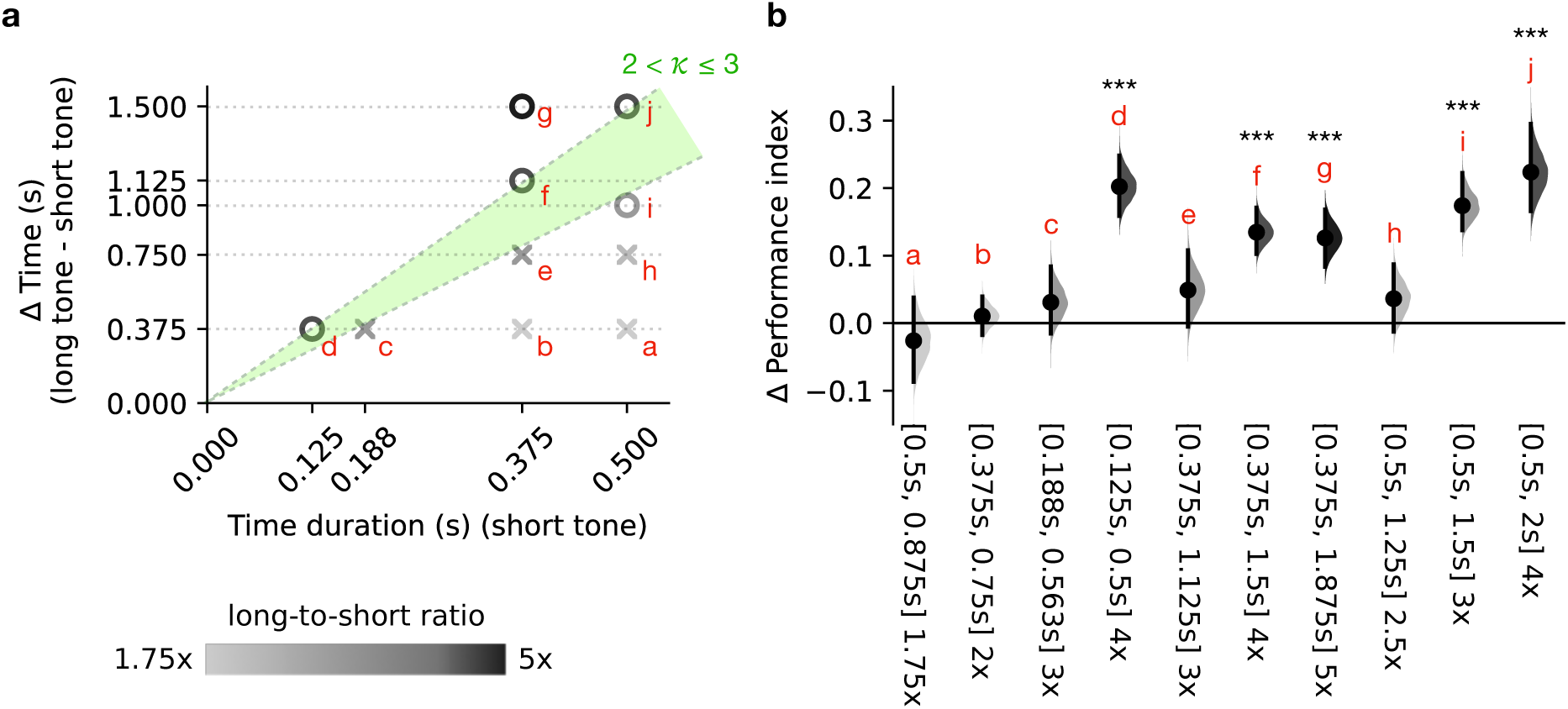
Temporal discrimination in flies follows Weber’s law. (a) Flies’ performance in the compound tone pattern learning task, in which they learn to associate a sugar reward with one of two compound tone patterns composed of short- and long-duration tones presented either in a short-to-long or long-to-short order. To successfully perform this associative learning task, flies must discriminate between the short-duration tone (x-axis) and the corresponding long-duration tone (x + ΔTime) within each pair. Each data point indicates success (⃝) or failure (×) of flies in learning the task with the indicated short/long tone-duration pair, reflecting their ability to discriminate the duration difference (ΔTime). Under Weber’s law, the Weber fraction (*k*) can be estimated by fitting a line through the origin, with the slope corresponding to *k*. Each line therefore represents a candidate discrimination threshold: tone-duration pairs above the line are predicted to be discriminable by flies, whereas those below the line are not. Because flies consistently discriminated tone pairs with a long-to-short duration ratio of 4 (points d, f, and j), but showed mixed performance at a ratio of 3 (non-discriminable at points c and e, but discriminable at point i), the data constrain the Weber fraction to a range rather than a single value. The green dashed lines define the boundaries of the shaded green area, which together indicate the range of plausible Weber fractions consistent with the observed behavioral data. Each data point (cross or circle) is shaded in grayscale according to the long-to-short duration ratio for the corresponding tone pair. (b) Data used to generate panel (a). Δ performance index represents the mean differences in performance between the Exp and Ctl groups, along with their 95% confidence intervals, for each tone-duration pair tested. The effect-size distributions are shaded using the same color code as in panel (a). The short and long tone durations used in each compound pattern are shown in brackets, followed by the ratio of long to short durations. *P*-values were calculated using Monte Carlo permutation test: ^∗∗∗^*P* < 0.001; no asterisk, *P* > 0.05.

According to Weber’s law, flies should detect a fixed time difference (ΔTime) more easily when the long-to-short duration ratio is higher, that is, when the short duration is smaller. Indeed, when ΔTime was kept at 0.375 s, flies failed to discriminate between the short and long tones when the short duration was 0.5 s (long/short ratio = 1.75), 0.375 s (ratio = 2), or 0.188 s (ratio = 3), but succeeded when it was 0.125 s (ratio = 4) (a, b, c, and d in Fig. 2). Notably, the performance index of flies showed an increasing trend as the short-duration decreased from 0.5 s to 0.188 s, followed by a sharp improvement in performance when it was further reduced to 0.125 s (ratio = 4) (a, b, c, and d in Fig. 2b). These results allow us to estimate the just-noticeable difference (JND), the minimum duration difference required for successful discrimination. Under Weber’s law, the ratio of the JND to the reference duration is known as the Weber fraction (*k*). Based on the observed discrimination threshold, the Weber fraction is estimated to be >2 and ≤3 (green area in Fig. 2a).

Next, we fixed the short duration at 0.375 s and tested whether flies exhibited improved performance as ΔTime increased. When ΔTime increased from 0.375 s (ratio = 2) to 0.75 s (ratio = 3), performance showed a slight but non-significant improvement (b and e in Fig. 2). A further increase to 1.125 s (ratio = 4) significantly enhanced performance, whereas no additional improvement was observed when ΔTime increased to 1.5 s (ratio = 5) (f and g in Fig. 2). These results further support the idea that flies’ ability to discriminate tone durations follows Weber’s law, with 2 < *k* ≤ 3. To further establish that flies’ interval discrimination follows Weber’s law, we tested additional long/short duration combinations. When the short duration was fixed at 0.5 s, flies failed to discriminate tone durations when ΔTime was 0.375 s or 0.75 s (ratios = 1.75 and 2.5, respectively), but succeeded when ΔTime increased to 1.5 s (ratio = 4) (a, h, j in Fig. 2). A slight deviation from Weber’s law was observed when ΔTime was 1 s (ratio = 3), where flies performed better than predicted based on the estimated Weber fraction (i in Fig. 2). Such deviations at higher stimulus magnitudes have been commonly reported in other sensory modalities and have been attributed to the saturation of neuronal variability as stimulus intensity increases^26^.

To further assess whether the Weber-like pattern in Fig. 2a could have arisen by chance, we performed a permutation test in which the success (⃝) and failure (×) labels were randomly reassigned across the tested short/long duration pairs (see Materials and Methods for details). For each permutation, we identified the Weberian boundary passing through the origin that best separated successful from failed discrimination. The observed data were significantly more consistent with such a boundary than expected by chance (*p* = 0.024, 100,000 permutations), supporting the conclusion that flies’ interval discrimination follows a proportional rule consistent with Weber’s law.

### Flies recognize relative temporal patterns across changes in absolute duration

The Weber-like pattern of interval discrimination highlighted the importance of duration ratios in temporal processing and was consistent with a ratio-based representation. Such a representation could support relative-timing generalization—the recognition of a learned temporal pattern even when its behaviorally salient elements are compressed or expanded, while their duration ratio is preserved. We therefore next asked whether flies show this capacity.

We first trained flies with compound patterns consisting of tones of longer absolute duration, either [0.5 s, 2 s] or [0.375 s, 1.5 s]; here, [A, B] denotes the two tone durations used to generate the short–long and long–short patterns. We then tested them with a novel and shorter [0.125 s, 0.5 s] pair. In both cases, the test pair preserved the same 1:4 duration ratio and short–long/long–short structure, while compressing the absolute tone durations fourfold or threefold. Flies showed a significant preference for the temporal pattern associated with reward during training in both tests, indicating that they could recognize the learned relative timing structure despite compression of the absolute tone duration (Fig. 3a and b). These results support relative-timing generalization in flies.

**Figure 3:**
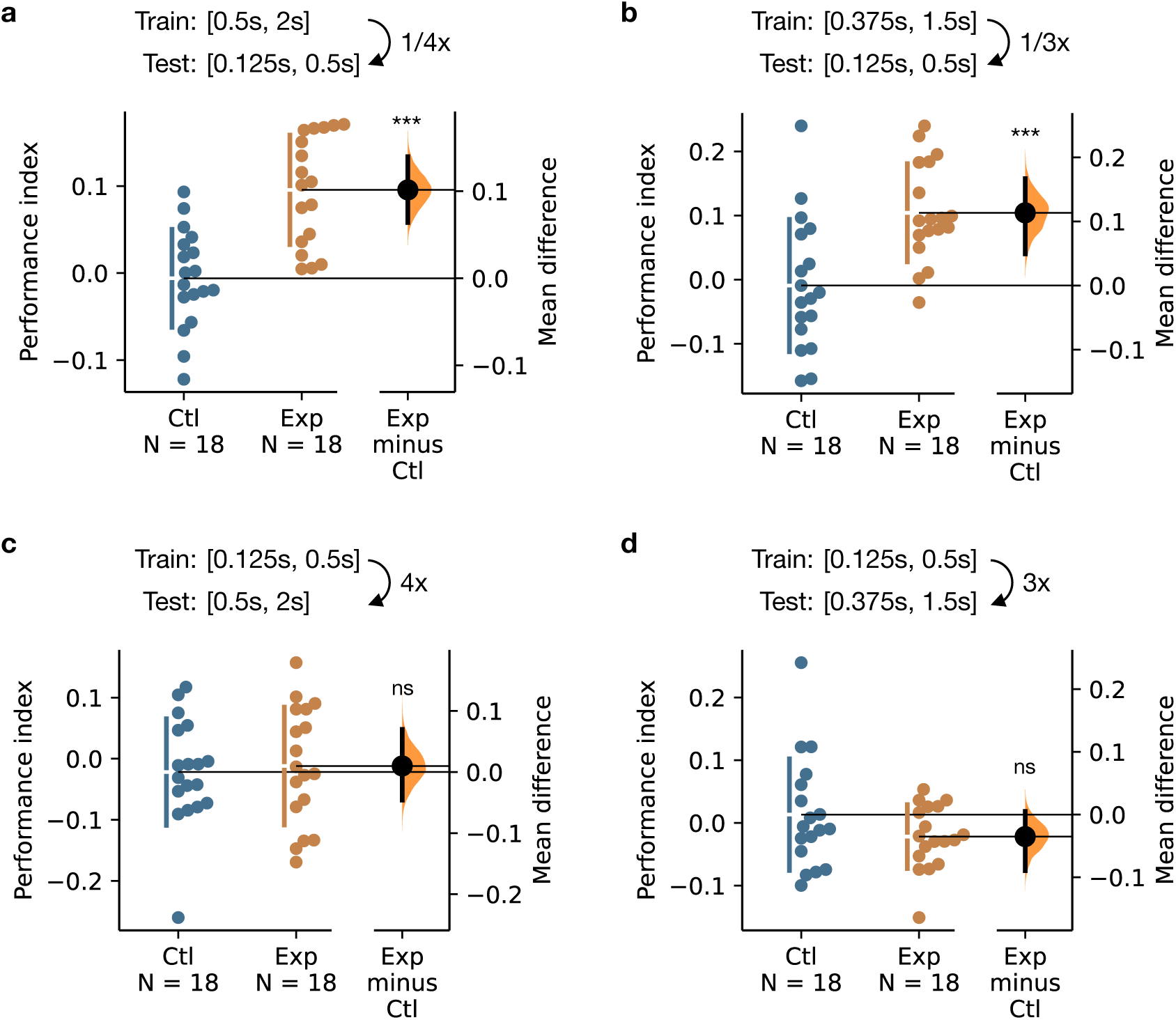
Flies generalize learned temporal patterns asymmetrically across changes in absolute duration. Flies trained with [0.5 s, 2 s] or [0.375 s, 1.5 s] compound tone patterns recognized the rewarded short–long or long–short structure when tested with the same structure proportionally compressed to [0.125 s, 0.5 s]. In contrast, flies trained with [0.125 s, 0.5 s] patterns failed to recognize the rewarded structure when it was proportionally expanded to [0.5 s, 2 s] or [0.375 s, 1.5 s]. (a) Trained with [0.5 s, 2 s] patterns and tested with [0.125 s, 0.5 s] patterns, corresponding to a fourfold compression (1/4x) of absolute tone durations. (b) Trained with [0.375 s, 1.5 s] patterns and tested with [0.125 s, 0.5 s] patterns, corresponding to a threefold compression (1/3x). (c) Trained with [0.125 s, 0.5 s] patterns and tested with [0.5 s, 2 s] patterns, corresponding to a fourfold expansion (4x). (d) Trained with [0.125 s, 0.5 s] patterns and tested with [0.375 s, 1.5 s] patterns, corresponding to a threefold expansion (3x). Data are presented as estimation plots showing mean differences between the Exp and Ctl groups, with bootstrap 95% confidence intervals shown on the right. *P*-values were calculated using Monte Carlo permutation test: ^∗∗∗^*P* < 0.001; ns, *P* > 0.05.

We next asked whether this generalization across changes in absolute duration was bidirectional. To test whether flies could also recognize the learned temporal pattern after temporal expansion, we trained them with the shorter [0.125 s, 0.5 s] pair and tested them with longer pairs, either [0.5 s, 2 s] or [0.375 s, 1.5 s]. In contrast to the compression experiments, flies failed to show a significant preference for the rewarded temporal pattern under either expansion condition (Fig. 3c and d). Thus, relative-timing generalization in flies appears to be asymmetric: learned relative timing structures can be transferred from longer to shorter durations, but not readily from shorter to longer durations.

This asymmetric generalization may reflect context-dependent calibration of temporal sensitivity^27–29^. Under Weberian timing, temporal discrimination becomes coarser as interval duration increases, such that longer intervals have a larger discrimination threshold than shorter intervals. Temporal representations may therefore be calibrated to the range of intervals experienced during training, with longer durations establishing a broader temporal comparison range than shorter durations. Such a broader range may readily accommodate compressed versions of the learned temporal structure, whereas a narrower range established by shorter durations may not extend to expanded test patterns. Together, these results show that flies recognize a learned temporal pattern from its relative timing structure, even as its elements are compressed.

### Flies generalize learned temporal patterns across sensory modalities

Weberian interval discrimination and relative-timing generalization indicate that flies encode temporal patterns flexibly. An especially stringent test of this idea is whether temporal information learned through one sensory modality can guide behavior through another. Success in such a task would imply that the fly brain can extract timing structure from the sensory signal that carries it, forming a representation that is not tied to a single modality. To test this possibility, we trained flies to associate auditory duration patterns with reward and then tested whether the same patterns, now presented as visual flashes rather than tones, could still guide them toward the rewarded choice.

Flies were first trained with [0.5 s, 2 s] compound tone patterns, as previously described. After training, they were presented with lights flashing in the same duration patterns as those used during training. Remarkably, both male and female flies showed a significant preference for the rewarded duration patterns, indicating that they can generalize temporal information learned in one modality to another (Fig. 4a, b). We further tested this capability using additional duration patterns. Flies were trained with [0.125 s, 0.5 s] compound tone patterns and tested with lights flashing in the same duration patterns. In this task, male but not female flies exhibited significant performance (Fig. 4c). We then attempted to make the task easier by increasing the difference between the short and long durations in the compound stimuli—from fourfold ([0.125 s, 0.5 s]) to fivefold ([0.1 s, 0.5 s]). However, even with this adjustment, only male flies exhibited significant performance (Fig. 4d). Thus, both sexes are capable of cross-modal temporal generalization, while males appear to retain this capacity across a broader range of temporal patterns.

**Figure 4:**
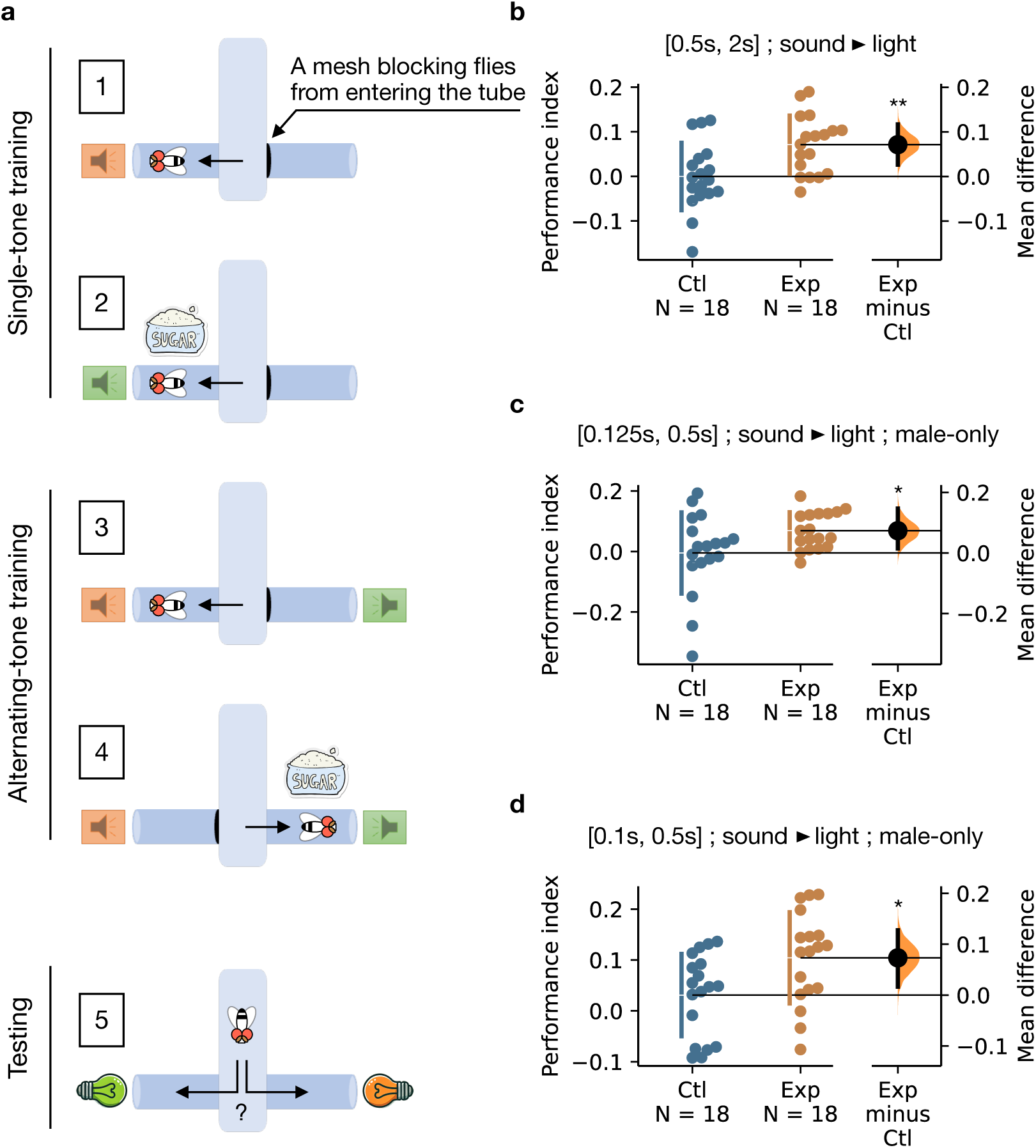
Learned temporal patterns can be generalized across sensory modalities. (a) Schematic of the training paradigm in which flies first learned to associate a sugar reward with compound tone patterns, then were tested with lights flashing in the same temporal patterns. Flies were presented with a pair of compound tone patterns (short–long vs. long–short), each lasting for 2 min. One pattern (CS–) was not paired with sugar (Step 1), while the other (CS+) was paired with sugar (Step 2). The flies were then presented with both tone patterns in alternation: first near the CS– (no sugar) for 2 min (Step 3), then near the CS+ (with sugar) for 2 min (Step 4). Immediately after training, the flies were tested for their preference between lights flashing in the same short–long and long–short duration patterns. The light cues alternated, and their sides were swapped to ensure that flies selected the rewarded pattern rather than a specific location (Step 5). The two duration patterns were counterbalanced as CS+, and performance scores were calculated as the average of two reciprocal training groups. (b) Flies trained with [0.5 s, 2 s] compound tone patterns preferred lights flashing in the same rewarded pattern, indicating generalization of temporal rules from the auditory to the visual modality. (c–d) Male flies showed similar preferences when trained with additional compound tone patterns: [0.125 s, 0.5 s] (c) and [0.1 s, 0.5 s] (d). The estimation plots in (b)–(d) show mean differences between the Exp and Ctl groups, with bootstrap 95% confidence intervals shown on the right. *P*-values were calculated using Monte Carlo permutation test: ^∗^*P* < 0.05, ^∗∗^*P* < 0.01.

The broader range of cross-modal generalization in males requires further investigation, but one possibility is that it reflects an adaptation associated with courtship behavior, where male flies must integrate multimodal cues to adjust the intricate temporal structure of their courtship song^30^. Such coupling of multimodal integration and temporal flexibility may underlie the enhanced generalization observed in males. Together, these results demonstrate that flies can transfer learned temporal patterns from one sensory modality to another, with males showing greater flexibility. This cross-modal transfer reveals a representation of time abstracted from the sensory channel that carried it.

### Compound tone pattern learning requires the mushroom body

To identify potential brain regions involved in processing and learning temporal information, we first focused on the MB. The MB is a major computational center in the fly brain, known to play key roles in associative learning, foraging, courtship, and sleep^31–34^. Its principal neurons, the Kenyon cells (KCs), receive diverse sensory inputs and transmit information to other brain regions via MB output neurons^35^. Although the role of the MB in auditory-guided behavior remains largely uncharacterized, it receives both direct and indirect input from neurons in the auditory pathway and is among the central brain regions that respond robustly to auditory stimuli^36^. Given its established roles in associative learning and multimodal sensory integration, the MB is a strong candidate for investigating how temporal information is processed and stored in the fly brain.

To test whether the MB is essential for compound tone pattern learning, we blocked synaptic output from the MB using the *UAS-shibire^ts^*^1^ (*UAS-shi^ts1^*) transgene^37^. This transgene encodes a temperature-sensitive dominant-negative form of dynamin that, when expressed in neurons, prevents synaptic vesicle recycling at the restrictive temperature (>29 °C), thereby blocking neurotransmission. We used *VT030559-GAL4* to drive *UAS-shi^ts1^* expression in all KCs and conducted compound tone learning experiments at 30 °C (Fig. 5a, b). Flies trained and tested with [0.5 s, 2 s] compound tone patterns exhibited significantly impaired performance at restrictive temperature of 30 °C (Fig. 5d), whereas their performance remained unaffected at the permissive temperature of 23 °C (Fig. 5c, e).

**Figure 5:**
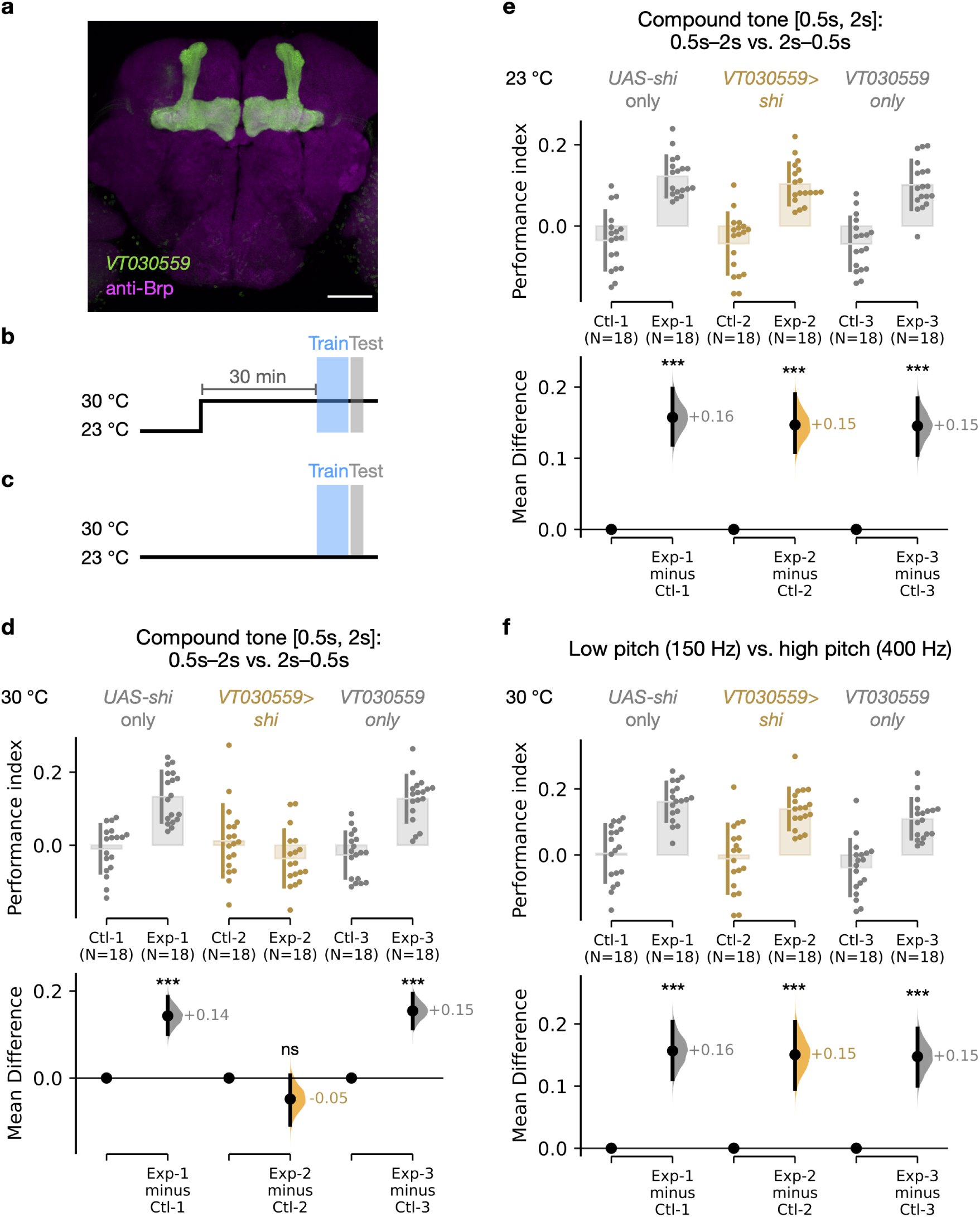
Compound tone pattern learning requires the mushroom body. (a) KCs labeled with mCD8::GFP driven by *VT030559-GAL4* (green). The brain was counterstained with anti-Brp antibody (magenta). Scale bar: 5 µm. (b-c) Temperature regime used for behavioral experiments. For restrictive condition (b), flies were raised and starved at 23 °C and were shifted to 30 °C 30 min before training and testing. For permissive temperature control (c), flies were raised, starved, trained, and tested entirely at 23 °C. (d) Flies failed to perform compound tone pattern learning (trained and tested with [0.5 s, 2 s] duration patterns) when KC neurotransmission was blocked by *shi^ts1^* at the restrictive temperature (30 °C). (e) Flies performed normally in compound tone pattern learning at the permissive temperature (23 °C). (f) Flies exhibited normal auditory pitch learning at the restrictive temperature (30 °C). The estimation plots in (d)–(f) show mean differences between the Exp and Ctl groups for each genotype, with bootstrap 95% confidence intervals displayed at the bottom. *P*-values were calculated using Monte Carlo permutation test: ^∗∗∗^*P* < 0.001; ns, *P* > 0.05.

To test whether the MB is specifically required for processing temporal information in this learning task, we examined whether blocking KC output affects other types of auditory learning that do not depend on tone-duration patterns. We trained flies using the same procedure as described in Fig. 1a, but replaced the compound tone patterns with continuous tones differing in pitch: a 150 Hz low-pitch tone (identical to the tone used in our previous experiments) and a 400 Hz high-pitch tone, which lies within the detectable range of the *Drosophila* auditory system^38^. During the single-tone training phase (Steps 1–2), each pitch tone was presented continuously for 2 min. During the alternating-tone phase (Steps 3–4), the two tones were played alternately from the opposite ends of the T-maze, each lasting for 3 s. The same alternating presentation was used during testing. Under this condition, flies were able to learn and discriminate between the pitch stimuli even when KC synaptic output was blocked by *shibire^ts1^* at the restrictive temperature, indicating that the MB is not required for auditory pitch learning (Fig. 5f). These results suggest that the role of the MB in auditory learning is specific to the processing of temporal patterns rather than general auditory cues. Together, these results identify the MB as a critical brain region for temporal pattern processing.

## Discussion

In this study, we demonstrate that flies can perceive and compare stimulus intervals in the subsecond-to-second range and use this temporal information to guide goal-directed behavior. Using a novel conditioning paradigm based on temporally patterned auditory cues, we show that flies can distinguish between tone-duration patterns and associate specific patterns with food rewards. This T-maze learning paradigm requires minimal behavioral shaping, making it well suited for probing flies’ innate capacity for interval discrimination. Because successful performance requires flies to distinguish between short- and long-duration tones, the assay provides a direct measure of duration discrimination. Notably, temporal discrimination depends on the ratio between durations rather than their absolute difference, consistent with Weber’s law, a fundamental principle in sensory perception. Beyond this proportional sensitivity, flies generalize learned temporal patterns across changes in absolute duration and across sensory modalities, suggesting that they form an abstract and modality-independent representation of time. Finally, we show that MB output is required for learning temporally patterned auditory cues.

Together, our study provides the first evidence that the fly brain supports ratio-based, generalizable, and cross-modal temporal computations. By showing that temporal flexibility follows Weber’s law, our findings demonstrate that a canonical psychophysical principle governs temporal computation even in a compact brain. They further extend recent evidence for flexible temporal cognition in eusocial insects^23^ to a non-eusocial, genetically tractable model. By demonstrating that MB output is required for temporal pattern learning, we identify a genetically accessible neural entry point for dissecting how such brains encode and use temporal information. Future studies investigating how the MB supports ratio-based interval comparison may provide key insights into the neural principles underlying temporal cognition.

### Weberian interval discrimination in *Drosophila*

In humans and many other vertebrates, interval discrimination often follows Weber’s law, with sensitivity governed by proportional rather than absolute differences in duration^3–9^. Whether this proportional rule also governs invertebrate interval discrimination, however, has remained unknown. In this study, we showed that *Drosophila* discriminates between tone durations based on their ratio rather than their absolute difference. Flies discriminated between durations when the longer interval was at least three to four times the length of the shorter one. In contrast, flies could not discriminate when the two durations differed by the same absolute amount but had a smaller ratio. These results indicate that the minimal time difference required for successful discrimination scales proportionally with the interval being timed, consistent with Weber’s law. The corresponding Weber fraction, which represents the slope of the proportional relationship, for time discrimination in flies is estimated to lie between 2 and 3. This proportional threshold falls within the range of stimulus ratios used in recent bumblebee studies, including the threefold long-to-short element ratio used to test abstract rhythm generalization^23^ and the fivefold duration difference used to test duration discrimination^22^. Thus, our findings place fly interval discrimination within an emerging comparative framework for studying how compact insect brains represent and compare time.

Flies’ performance in duration discrimination showed a slight deviation from Weber’s law when the tone durations reached the longer end of the tested range (i in Fig. 2). This pattern echoes observations in other sensory systems^26,39–41^. Such departures from Weber’s law have been linked to the saturation of neuronal variability at higher stimulus magnitudes^26^. Our findings suggest that similar neural constraints may also shape temporal processing in *Drosophila*. Taken together, these results suggest that Weberian temporal processing—implemented, for example, through accumulator mechanisms with interval-scaled variability or logarithmically compressed magnitude codes^1,13,42,43^—can also operate in a compact invertebrate brain, whether inherited from a common ancestor or independently evolved through convergent evolution.

### Generalizable temporal representations in *Drosophila*

By encoding intervals proportionally, a Weberian representation provides a potential means for an animal to recognize temporal structure despite changes in the absolute durations through which that structure is expressed. In our experiments, flies trained to associate reward with one temporal pattern (short–long or long–short) could later choose the corresponding pattern when the absolute tone durations were compressed, while their order and duration ratio were preserved. This indicates that their behavior was not governed solely by memorized auditory templates or fixed interval values. Rather, flies retained information about the relationship between intervals, allowing a learned temporal rule to guide behavior under a new timing scale.

Interestingly, the same temporal relationship did not generalize when the absolute durations were expanded. One possible explanation for this asymmetry is that temporal representations may be calibrated to the range of intervals experienced during training. Sensory systems often adapt their coding range to recent stimulus statistics, allowing perception and behavior to remain sensitive within the most relevant range. For example, adaptive coding in the fly visual system rescales neural responses to changing stimulus statistics^27^, auditory neurons adjust their dynamic range to sound-level distributions^28^, and *Drosophila* larvae adapt navigational decisions to stimulus variance^29^. A similar principle may apply to temporal learning. Because the absolute just-noticeable difference between intervals grows with their duration under Weberian timing^1,2,6,7^, the temporal comparison range can expand with interval duration rather than maintaining a fixed absolute resolution. Flies trained with longer duration pairs may establish a broader temporal comparison range than flies trained with shorter pairs. As a result, training with longer intervals may permit generalization to compressed test patterns, whereas training with shorter intervals may not support extrapolation to expanded patterns.

Beyond relative-timing generalization, we also found that flies can transfer learned temporal relationships across sensory modalities. Cross-modal transfer of temporal learning has been reported in several vertebrates, including rats, pigeons, and rabbits, which can apply learned temporal rules across auditory, visual, and somatosensory cues^44–49^. The same is true in humans, where cross-modal transfer of temporal discrimination points to a shared, modality-independent timing system^50–53^. Recent work in bumblebees showed that learned rhythmic patterns can transfer from vibration to vision, indicating that cross-modal temporal representations can also occur in insects^23^. Our findings extend this principle to *Drosophila*, showing that a non-eusocial insect can transfer learned interval-duration rules from auditory to visual cues. This finding suggests that such temporal representations are not restricted to animals with the complex social behaviors thought to place particularly high demands on temporal cognition. Together, these findings suggest that modality-independent temporal representations may be broadly accessible to compact brains and can emerge across insects with distinct ecological demands and neural architectures.

### The MB and time representations in flies

The MB is required for flies to perform successfully in our temporal learning paradigm. When KC output was selectively blocked using temperature-sensitive *shibire^ts1^* during training and testing, flies failed to show a preference for the rewarded tone patterns. The MB is best known for its role in associative learning, where sensory inputs and reinforcement signals converge to drive conditioned behavior^34,54–57^. However, the MB is not required for all types of learning. For example, flies can successfully perform spatial learning tasks, such as locating a cool spot using visual cues, even when MB function is disrupted^58^. The MB’s requirement in our temporal learning task suggests that representations of tone-duration patterns are likely formed or accessed within this structure.

This interpretation is strengthened by a recent study showing that the MB can support interval-specific olfactory learning^21^. In that work, individual KCs reached peak activity at distinct, odor-specific delays after odor onset, such that the KC population collectively formed a distributed representation of elapsed time. Combined with dopamine-gated synaptic plasticity, this sequential KC activity provides a circuit mechanism by which flies could learn to associate reinforcement with a particular interval following a sensory cue. These findings therefore provide important independent support for the idea that the MB is capable of encoding temporal information. At the same time, the framework based on sequential KC activity and dopamine-gated plasticity does not readily explain the more flexible timing abilities revealed by our task, including ratio-based discrimination of complex tone-duration patterns and generalization of temporal rules across changes in absolute duration and sensory modality. Taken together, however, these findings position the MB as a promising entry point for dissecting temporal computation in a compact, genetically accessible brain, with the potential to reveal general principles of time processing across species.

## Acknowledgements

We thank Wei-Ming Fu, Wen-Chun Lo, Hung-Tse Lin, Pei-Hsun Hsieh, Zhenjiang Chen, and Tristan Hsu for carrying out initial studies of the present work. S. Lin is supported by the Academia Sinica Grand Challenge Seed Grant (GBA-107-TP-115-06), Academia Sinica Career Development Award (AS-CDA-110-L06), and a grant from Taiwan National Science and Technology Council (113-2628-B-001-009-MY3). C.-Y. Huang is supported by the Academia Sinica Scholar Award (ASSA-113-04) and grants from Taiwan National Science and Technology Council (107-2410-H-002-020-MY3 and 110-2410-H-002-212-MY3).

## Author contributions

T.-N.L. performed all experiments and contributed to study design, data analysis and interpretation. C.-Y.H. contributed to study design, data analysis and interpretation, and revised the manuscript. S.L. contributed to study design, data analysis and interpretation, and drafted the manuscript. C.-Y.H. and S.L. co-supervised the study.

## Materials and methods

### Fly husbandry

Flies (*Drosophila melanogaster*) were raised on standard fly food on a 12 h–12 h light–dark cycle at 23 °C with 50% humidity. Flies used in all the experiments in this study are 5∼7 days old adults. The following fly lines were used: Canton-S, *VT030559-GAL4* (VDRC: 206077), and *UAS-shibire^ts1^* (Kitamoto, 2001)^37^.

### Temporal learning paradigms

Training and testing were conducted in a T-maze apparatus (CelExplorer Labs Co., TM-101). Flies were starved for 24 h prior to training to enhance their motivation for sugar-seeking. During tone pattern training, flies were presented with two distinct tone patterns composed of long- and short-duration tones. One pattern (CS–) was presented without sugar, while the other pattern (CS+) was paired with sugar. Each tone pattern was presented continuously for 2 min during its respective training phase. Flies were then exposed to both patterns played in alternation. They were first confined to the arm near the CS– pattern (no sugar) for 2 min, and subsequently confined to the arm near the CS+ pattern (with sugar) for another 2 min. After training, the locations of the two tone patterns were swapped, and flies were immediately tested for their preference between the CS+ and CS– patterns, which were again presented in alternation. To control for innate preferences, CS+ and CS– assignments were counterbalanced across groups, and a performance index was calculated as the average of the two reciprocal training groups.

In all experiments, flies were assigned to an experimental group (Exp), in which the CS+ pattern was paired with sugar, or to a control group (Ctl), which underwent the same training procedure without sugar. Flies were considered to have successfully learned the task when the Exp group performed significantly better than the Ctl group. Male and female flies were trained and tested together, and the flies in each arm of the T-maze were counted separately by sex. Except in Fig. 4c and d, performance indices were calculated from the combined counts of both sexes; Fig. 4c and d show male performance indices only.

Tone duration patterns included either simple repeated tones with fixed durations or compound sequences combining short- and long-duration tones. In simple patterns, a single tone duration was repeated throughout training, with the silent gap between tones adjusted so that the total number of tones presented during training was the same across patterns of different tone durations. In compound patterns, each cycle consisted of a short- and a long-duration tone separated by a 0.5-s silent gap (in short–long or long–short order), followed by a 3-s silent gap before the next cycle began. Detailed configurations of the tone patterns are described in the main text. Tone patterns were generated using GarageBand (Apple Inc.) and played through a Mi Portable Bluetooth Speaker (XMYX04WM). All tones were 150 Hz sine waves played at 90 dB, measured using a decibel meter (MET-SLM1358) (Supplementary Materials 1 and 2).

For cross-modal learning experiments, flies were trained with compound tone patterns and tested with lights flashing in the same duration patterns. For pitch discrimination experiments, we used the same training and testing procedure as described above, but replaced the temporal tone patterns with two continuous tones of distinct pitch: 150 Hz (low) and 400 Hz (high) (Supplementary Materials 3 and 4). During the single-tone training phase (Steps 1–2), each tone was played continuously for 2 min. During the alternating-tone training phase (Steps 3–4), the two tones were played alternately from opposite arms of the T-maze in 3-s repeats (i.e., 3 s of low pitch followed by 3 s of high pitch). The same alternating presentation was used during testing. These stimuli were presented at 90 dB.

### Permutation test for figure 2a

To assess whether the observed alignment between behavioral outcomes and a Weber-fraction line could occur by chance, we performed a permutation test based on label shuffling. The coordinates of the ten tested compound tone pairs (short tone duration on the x-axis and ΔTime on the y-axis) were fixed, and the five success (⃝) and five failure (×) labels were randomly reassigned across these positions. For each permutation, we identified the line through the origin that best separated the shuffled labels. This line corresponds to the best-fitting Weber fraction, with successes expected to fall on or above the line (i.e., ΔTime is equal to or greater than the JND) and failures below it (i.e., ΔTime is smaller than the JND). Any violation of this rule was counted as a misclassification, and the total number of violations defined the misclassification score for the permutation. In the observed data, the best-fitting line yielded a score of 1 (only point i was misclassified). To evaluate the likelihood of obtaining such a low score by chance, we generated a null distribution by repeating the label-shuffling procedure 100,000 times and recording the best-fitting misclassification score from each permutation. The empirical *p*-value was then calculated as the proportion of permutations whose score was less than or equal to the observed value. The Python script used for this analysis is provided in the Supplementary Materials (Supplementary Materials 5).

### Immunostaining

Fly brains were dissected in PBS (Sigma, P4417) and fixed in PBS containing 4% formaldehyde (Sigma, F8775) for 20 min at room temperature. After fixation, brains were washed three times in PBST (0.5% Triton X-100 in PBS) for 20 min each, then blocked in PBST containing 5% normal goat serum (Jackson ImmunoResearch, RRID:AB_2336990) for 30 min. Brains were then incubated with primary antibodies in PBST containing 5% normal goat serum at 4 °C overnight. The following day, the brains were washed three times in PBST for 20 min each at room temperature, followed by incubation with secondary antibodies in PBST at 4 °C overnight. Finally, brains were washed three times in PBST for 20 min each at room temperature and mounted with Gold Antifade reagent (Thermo Fisher Scientific, S36937). The primary antibodies used were mouse anti-Brp (1:100; DSHB, RRID:AB_2314866) and rat anti-mCD8*α* (1:100; Thermo Fisher Scientific, RRID:AB_10392843). The secondary antibodies were goat anti-mouse (Cy3) (1:400; Jackson ImmunoResearch, RRID:AB_2338692) and goat anti-rat (Alexa 488) (1:400; Jackson ImmunoResearch, RRID:AB_141373).

### Statistical analysis

Statistical analyses and estimation plots were generated using the Python package DABEST^59^. Effect sizes (mean differences between the Exp and Ctl groups) were reported together with bootstrap 95% confidence intervals, which are robust to non-normal data. *P*-values were obtained with a nonparametric, two-sided approximate permutation *t* test (Monte Carlo permutation test) using 5,000 permutations, and were adjusted according to Phipson and Smyth (2010)^60^.

### Use of AI tools

ChatGPT 5.5 (OpenAI) was used to help write the Python script for the permutation test (Supplementary Materials 5); the authors verified the code and its output.

### Data availability

All behavioral data, imaging data and analysis code underlying the findings of this study will be deposited in a public repository upon acceptance, and the accession details will be provided in the final version of the article.

## Supplementary Materials

**Supplementary Materials 1. Short–long pattern in the [0.5 s, 2 s] compound tone.** Repeated short–long sequence used to construct the [0.5 s, 2 s] compound tone.

**Supplementary Materials 2. Long–short pattern in the [0.5 s, 2 s] compound tone.** Repeated long–short sequence used to construct the [0.5 s, 2 s] compound tone.

**Supplementary Materials 3. Low-pitch tone (150 Hz).** 150-Hz tone used in Fig. 5f.

**Supplementary Materials 4. High-pitch tone (400 Hz).** 400-Hz tone used in Fig. 5f.

**Supplementary Materials 5. Permutation test code for Fig. 2a**. Python script used to run the permutation test for Fig. 2a.

